# C_4_-dicarboxylate metabolons: Interaction of C_4_-dicarboxylate transporters of *Escherichia coli* with cytosolic enzymes

**DOI:** 10.1101/2021.03.01.433382

**Authors:** Christopher Schubert, Gottfried Unden

## Abstract

Metabolons represent the structural organization of proteins for metabolic or regulatory pathways. Here the interaction of enzymes fumarase FumB and aspartase AspA with the C4-DC transporters DcuA and DcuB of *Escherichia coli* was tested by a bacterial two-hybrid (BACTH) assay *in situ*, or by co-chromatography (mSPINE). DcuB interacted strongly with FumB and AspA, and DcuA with AspA. The *fumB-dcuB* and the *dcuA-aspA* genes encoding the respective proteins are known for their colocalization on the genome and the production of co-transcripts. The data consistently suggest the formation of DcuB/FumB, DcuB/AspA and DcuA/AspA metabolons in fumarate respiration for the uptake of L-malate, or L-aspartate, conversion to fumarate and excretion of succinate after reduction. The DcuA/AspA metabolon catalyzes L-Asp uptake and fumarate excretion in concerted action also to provide ammonia for nitrogen assimilation.

## Introduction

Versatile bacteria such as *Escherichia coli* co-ordinate metabolism and cellular physiology on various levels including gene expression and post-translational regulation for optimal adaption to current growth conditions. For stabilization and improvement of linear pathways, the enzymes and proteins can be organized in metabolons. Metabolons represent weak complexes of enzymes catalyzing consecutive reactions in order to transfer intermediates between the enzymes of pathways. Channeling prevents the release of labile or toxic intermediates, supports higher local concentration of the intermediates and prevents drifting to other pathways. This organization is able to increase metabolic efficiency and provide an opportunity to control the flux. High-throughput approaches have been used to identify complexes of weakly interacting proteins, or ‘interactomes’, in microorganisms, including proteins linking membrane processes to cytoplasmic reactions (Butland *et al.* 2005; Hu *et al.* 2009). However, the general approaches provide limited information on complexes when partner proteins are present in low concentrations or are membrane-bound. Thus, established regulatory complexes of the membrane-integral C4-dicarboxylic acids (C4-DC) sensor kinase DcuS with the coregulatory C4-DC carriers DctA, DcuB or DauA (Kleefeld *et al.* 2009b; Witan *et al.* 2012; Karinou *et al.* 2013; Unden *et al.* 2016) and others were not identified. Moraes and Reithmeier (2012) suggested an alternative approach for identifying gene clusters of *E. coli* for metabolons consisting of membrane integral transporters together with cytosolic enzymes for the subsequent pathway. The overview Moraes and Reithmeier (2012) lists polycistronic operons encoding on the one hand transporters for the uptake of sugars, sugar derivatives, amino acids, C4-DC, and other substrates, and on the other hand, enzymes or regulators for subsequent pathways. For some systems interaction of the proteins has been demonstrated, such as for AmtB-GlnK (ammonia transporter and Pll-type regulator GlnK) (van Heeswijk *et al.* 1996; Coutts *et al.* 2002). Remarkably, for most of the genetically co-localized systems interaction of the transporter/enzyme pairs is hypothetical.

The list of potential metabolons (Moraes and Reithmeier 2012) also includes the operons *fumB-dcuB* encoding fumarase FumB and the C4-dicarboxylate transporters DcuB, *dcuA-aspA* encoding the C4-dicarboxylate transporters DcuA and aspartase AspA (Six *et al.* 1994; Golby *et al.* 1998b), and *ttdAB-ttdT* encoding tartrate dehydratase TtdAB and the L-tartrate transporter TtdT (Kim and Unden 2007). Both *dcuA* and *dcuB* are transcribed independently from *fumB* and *aspA,* respectively, but also as *fumB-dcuB* and *dcuA-aspA* cotranscripts (Golby *et al.* 1998b). DcuB is produced only under anaerobic conditions (Zientz *et al.* 1998; Golby *et al.* 1999) and catalyzes the uptake of L-malate which is then dehydrated by FumB to fumarate (Engel *et al.* 1994; Six *et al.* 1994) (Fig. 1A). Fumarate is reduced to succinate by fumarate reductase in fumarate respiration. Succinate is not further catabolized anaerobically and excreted by DcuB via a L-malate/succinate antiport. DcuB also catalyzes the uptake of fumarate which is used directly in fumarate respiration. In the same way, L-Asp is a substrate for transport by DcuB and used after deamination by AspA to fumarate for fumarate respiration. In any case, the succinate is excreted by C4-DC/succinate antiport with C4-DC indicating any of the C4-DCs L-malate, fumarate, or L-aspartate (Engel *et al.* 1994; Six *et al.* 1994; Janausch *et al.* 2002; Schubert *et al.* 2021).

**Fig. 1:**
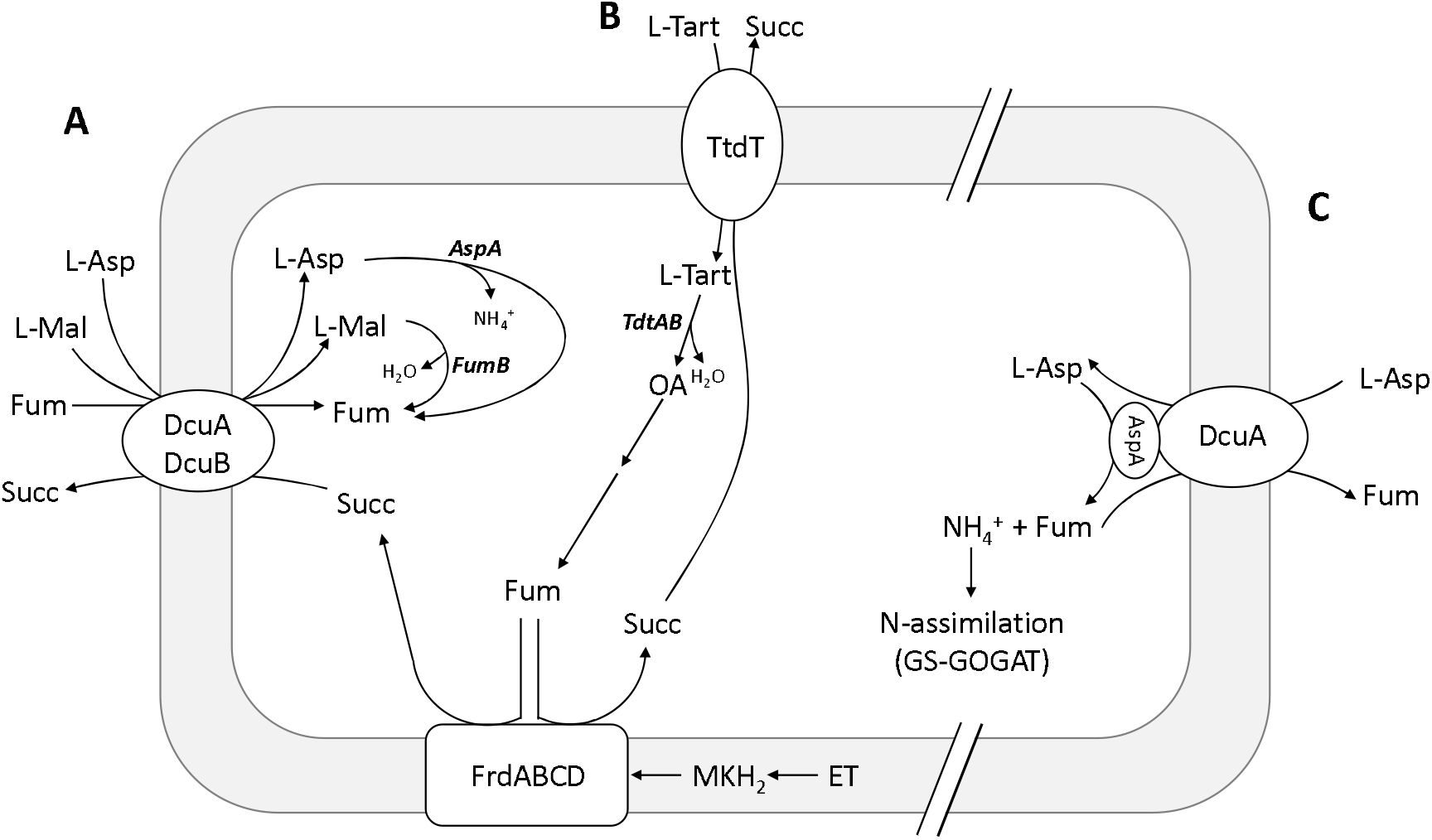
Pathways of (A) fumarate respiration of *E. coli* using fumarate, L-malate, and L-aspartate as the electron acceptors, (B) L-tartrate fermentation and (C) nitrogen assimilation from L-aspartate. For details on (A) and (B) see Unden *et al.* (2016) and on (C) Schubert *et al.* (2020). Abbreviations: AspA, aspartase, DcuA, C4-DC transporter DcuA; DcuB, C4-DC transporter DcuB, ET, electron transport; FrdABCD, fumarate reductase; Fum, fumarate; FumB, fumarase B; L-Asp, L-aspartate; L-Mal, L-malate; MKH_2_, menaquinol; Succ, succinate; TtdT, L-tartrate transporter.

DcuA on the other hand is expressed constitutively under aerobic and anaerobic conditions (Golby *et al.* 1998b). DcuA is able to substitute DcuB in anaerobic growth (Six *et al.* 1994) and catalyze an C4-DC/succinate antiport for fumarate respiration (Fig. 1A). L-Asp, fumarate or L-malate are accepted as the C4-DCs, but with a clear preference for L-Asp over L-malate or fumarate (Strecker *et al.* 2018). In addition to its role in catabolism and fumarate respiration, DcuA catalyzes the uptake of L-aspartate for anabolism (Strecker *et al.* 2018; Schubert *et al.* 2020). Under these conditions (Fig. 1B) L-aspartate is deaminated in bacteria by AspA to fumarate and ammonia, which can saturate the complete nitrogen requirement for growth.

The uptake of L-Asp depends on DcuA and is coupled to the excretion of fumarate (and some L-malate). The combined action of DcuA and AspA results in the net uptake of ammonia, which is able to supply all the nitrogen for cell synthesis without consuming the carbon, provided that other carbon sources are available.

L-Tartrate that is taken up by TtdT for tartrate fermentation (Fig. 1A), has to be dehydrated by dehydratase TtdAB to oxaloacetate (Reaney *et al.* 1993; Kim and Unden 2007). After conversion to fumarate, it is used in fumarate respiration, and the succinate is excreted by

### TtdT in antiport against L-tartrate

In the metabolic pathways, DcuA, DcuB and TtdT are closely linked to AspA in case of DcuA and DcuB, to FumB in the case of DcuB, and to TtdAB in the case of TtdT. To test the metabolon hypothesis for the C4-dicarboxylate metabolism, interaction of DcuA and DcuB with AspA and FumB was tested. Interaction was tested *in vivo* by the adenylate cyclase-based bacterial two-hybrid (BACTH) system which has been established as a reliable assay for protein-protein interactions involving membrane-integral proteins, or *in vitro* by copurification of the proteins.

## Results and Discussion

### DcuB interacts with AspA and FumB

The adenylate cyclase-based bacterial two hybrid (BACTH) system (Karimova *et al.* 1998; Karimova *et al.* 2001) was used to investigate the interaction of AspA or of FumB with DcuB. The BACTH system relies on the restoration of *Bordetella pertussis* adenylate cyclase (AC) that has been separated genetically in two domains T18 and T25. When the domains are fused to interacting proteins, local vicinity of the domains allows restoration of AC activity due to high flexibility of domain arrangement of *B. pertussis* AC. The activity is tested in a reporter strain deficient of the *E. coli* AC CyaA by expression of the cAMP-CRP dependent β-galactosidase LacZ. The N- and C-termini of DcuB (similar to those of DcuA and DcuC) are located in the periplasm, which excludes the use of the BACTH system under standard conditions since cAMP-CRP resides in the cytoplasm. Therefore, the T25 fragment of the AC was fused for the interaction assays into a cytosolic loop of DcuB (or the other Dcu transporters) (Bauer *et al.* 2011; Witan *et al.* 2012; Wörner *et al.* 2016; Strecker *et al.* 2018). DcuB with the sandwich (SW) T25 domain (DcuB_SWT25_) is membrane-integral and active in coregulating DcuS (Wörner *et al.* 2016) allowing interaction studies. Bacteria producing the pairs DcuB_SWT25_ with FumB_T18_, and DcuB_SWT25_ with AspA_T18_ showed high β-galactosidase activity (Fig. 2) which exceeded the negative control represented by the non-interacting proteins _T18_AspA and _T25_Zip. The interaction was independent of the presence of transport substrate molecules L-malate and L-aspartate (not shown). On the other hand, bacteria producing the sandwich construct DcuC_SWT25_ and FumB_T18_, or AspA_T18_ respectively, showed only low β-galactosidase activity, which exceeded the background activity only slightly (Fig. 2). DcuC serves as a C4-DC efflux transporter (Zientz *et al.* 1996; Zientz *et al.* 1999; Janausch *et al.* 2002) for succinate derived from fermentation metabolism and not from externally supplied C4-DCs. The data indicates specific protein-protein interaction of DcuB with FumB and with AspA which is not observed for the DcuC efflux transporter that has a different metabolic task.

**Fig. 2:**
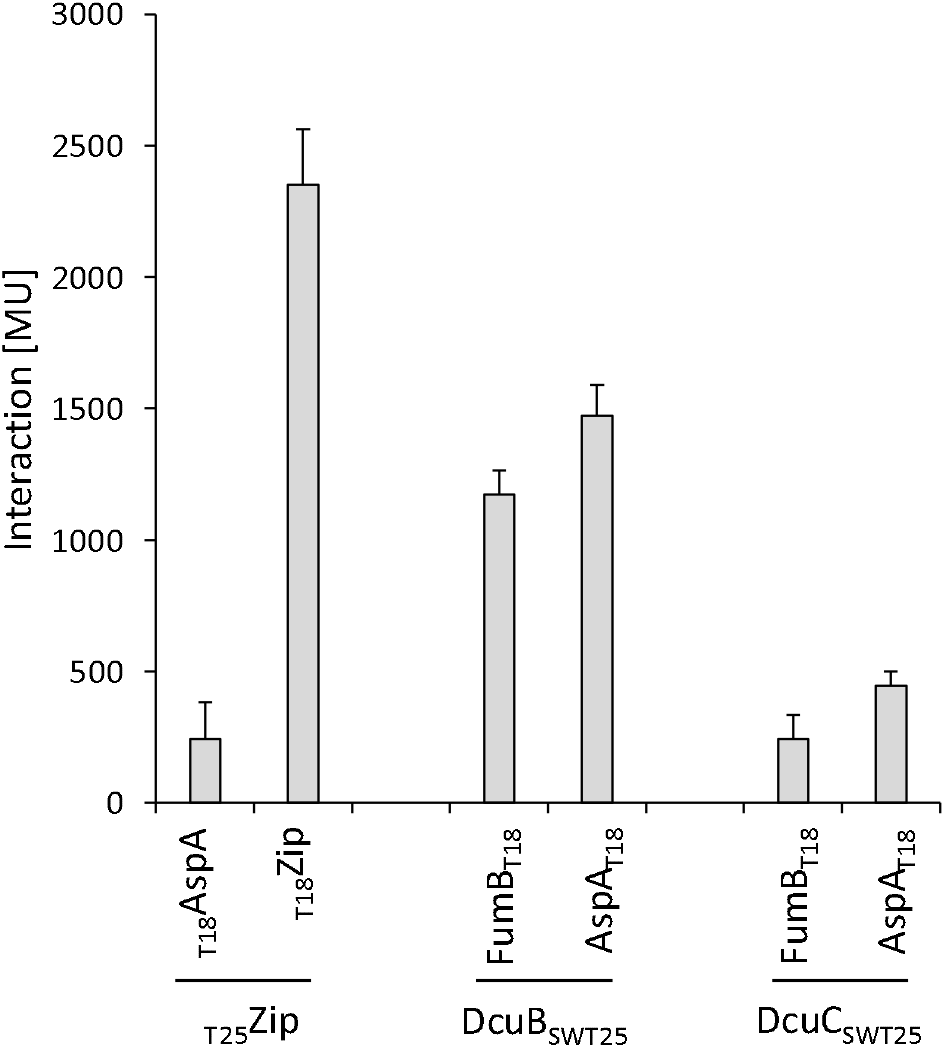
Interaction of DcuB and DcuC with AspA and FumB tested in the BACTH system. *E. coli* BTH101(Δ*cyaA*) was co-transformed pairwise with plasmids encoding T25 sandwich constructs with DcuB (DcuB_SWT25_) or DcuC (DcuC_SWT25_) and fusions of FumB or AspA to T18 (FumB_T18_, and AspA_T18_). The combinations are shown on the x-axis. The leucine zipper pairs _T18_Zip and _T25_Zip are applied as positive control (Karimova *et al.*, 1998; 2001). The pair _T18_Zip/_T25_AspA represents a negative control for background β-galactosidase activity. The corresponding plasmids are derivatives of pUT18 (FumB_T18 and_ AspA_T18_) and pKNT25 (DcuB_SWT25_) (Table 1). Bacteria were grown anaerobically in LB medium, β-Galactosidase activities were quantified in Miller-Units MU (Miller 1992).

### DcuA interacts with AspA

The role of DcuA in L-Asp uptake for fumarate respiration in catabolism, and as a source of nitrogen suggests a close metabolic link to aspartase AspA. C- and N-termini of DcuA are located in the periplasm (Golby *et al.* 1998a) which prevents their use for fusion to the T18 and T25 AC domains for the BACTH system. Insertion of the T25 domain into a cytosolic loop of DcuA transporter in a sandwich fusion (DcuA_SWT25_) produced a fusion protein in analogy to DcuB_SWT25_. The fusion protein was located in the membrane but was not active in transport (Strecker *et al.* 2018) or any other biological activity, which makes its application in the BACTH assay questionable. In an alternative approach, DcuA was fused to the bacterial alkaline phosphatase PhoA. PhoA-tagged membrane protein like DcuB often retain biological activity (Bauer *et al.* 2011) and the tag can be used for immunodetection with PhoA antisera. In the experiment, *E. coli* strain C43 was co-transformed with plasmids encoding AspA-Strep and DcuA-PhoA fusions. Proteins were simultaneously overexpressed and the bacteria were treated with formaldehyde after expression to crosslink proteins that are located in close proximity (Sutherland *et al.* 2008). Detergent extracts of the membranes were then applied to a Strep-Tactin column. After a wash step, the specifically bound proteins were eluted with desthiobiotin. A column treated with the extracts from bacteria that produced DcuA-PhoA and AspA-Strep eluted DcuA-PhoA and AspA-Strep from the column upon addition of desthiobiotin (Fig. 3). DcuA-PhoA was missing, however, in the eluate when either AspA-Strep or DcuA-PhoA were not produced by the bacteria. The experiment demonstrates that interaction with AspA-Strep is required to retard DcuA to the Strep-Tactin column. In addition, binding of DcuA to AspA-Strep depended on formaldehyde cross-linking in the cell homogenate (not shown), supporting the notion that the interaction is too weak to survive column chromatography without stabilization by crosslinking.

**Fig. 3:**
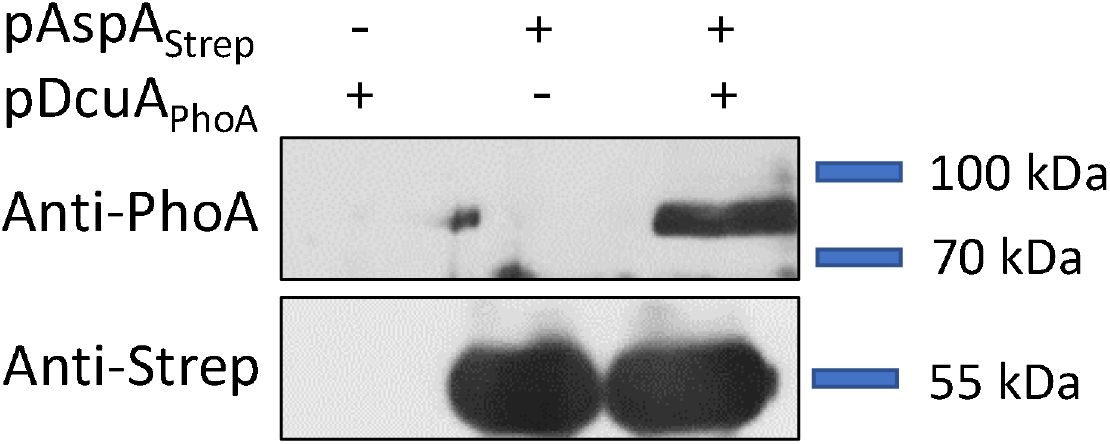
Copurification of DcuA-PhoA with AspA-Strep on a Strep-Tactin column. Strep-tagged AspA (AspA-Strep, encoded by pMW3049) and PhoA-tagged DcuA (DcuA-PhoA, encoded by pMW577) were expressed individually or co-expressed in *E. coli* C43. For protein crosslinking, bacterial cell suspensions were incubated with formaldehyde (0.6 % w/v) for 15 min: Then the bacteria were homogenized in a French press cell and treated with lauryldimethylamine oxide (0.05% v/v). The cleared cell homogenate was applied to a Strep-Tactin column. After washing, the bound proteins were eluted with buffer containing desthiobiotin (2.5 mM). Samples were subjected to SDS-PAGE and Western blotting. Western blots were probed with antibodies against PhoA (upper panel) and Strep-tag (lower panel). Protein standards are shown on the right-hand side. The masses of AspA-Strep and DcuA-PhoA calculated from amino acid composition are 56.0 kDa and 95.2 kDa, respectively.

### Function of DcuA/AspA, DcuB/FumB and DcuB/AspA metabolons?

The experiments show specific interaction of the C4-DC transporters DcuA with AspA, and of DcuB with FumB and AspA (Fig. 4). Under anaerobic conditions fumarate represents the substrate for fumarate respiration. DcuB is a major carrier for C4-DC/succinate antiport in fumarate respiration (Six *et al.* 1994; Golby *et al.* 1998b; Janausch *et al.* 2002; Unden *et al.* 2016). The major substrate for DcuB is L-malate, but due to the high levels of L-aspartate in the intestine of animals (Bertin *et al.* 2018; Schubert *et al.* 2021), L-Asp is also transported by DcuB (Six *et al.* 1994; Golby *et al.* 1998b). The interaction of DcuB with AspA and FumB suggests a coordination of the C4-DC uptake with the conversion to fumarate, and the excretion of the succinate after fumarate reduction, which indicates in the presence of DcuB/FumB and DcuB/AspA metabolons (Fig. 4A). A very similar situation applies to the role for a DcuA/AspA metabolon in fumarate respiration (Fig.4A). Overall, DcuB/FumB, DcuB/AspA and DcuA/AspA metabolons are consistently indicated for fumarate respiration by interaction of the proteins, the colocalization of the corresponding genes in the genome and the physiological relevance of a coordination (Fig. 4A). The metabolons await, however, direct proof of metabolic coupling and biochemical backgrounds of the coupling.

**Fig. 4:**
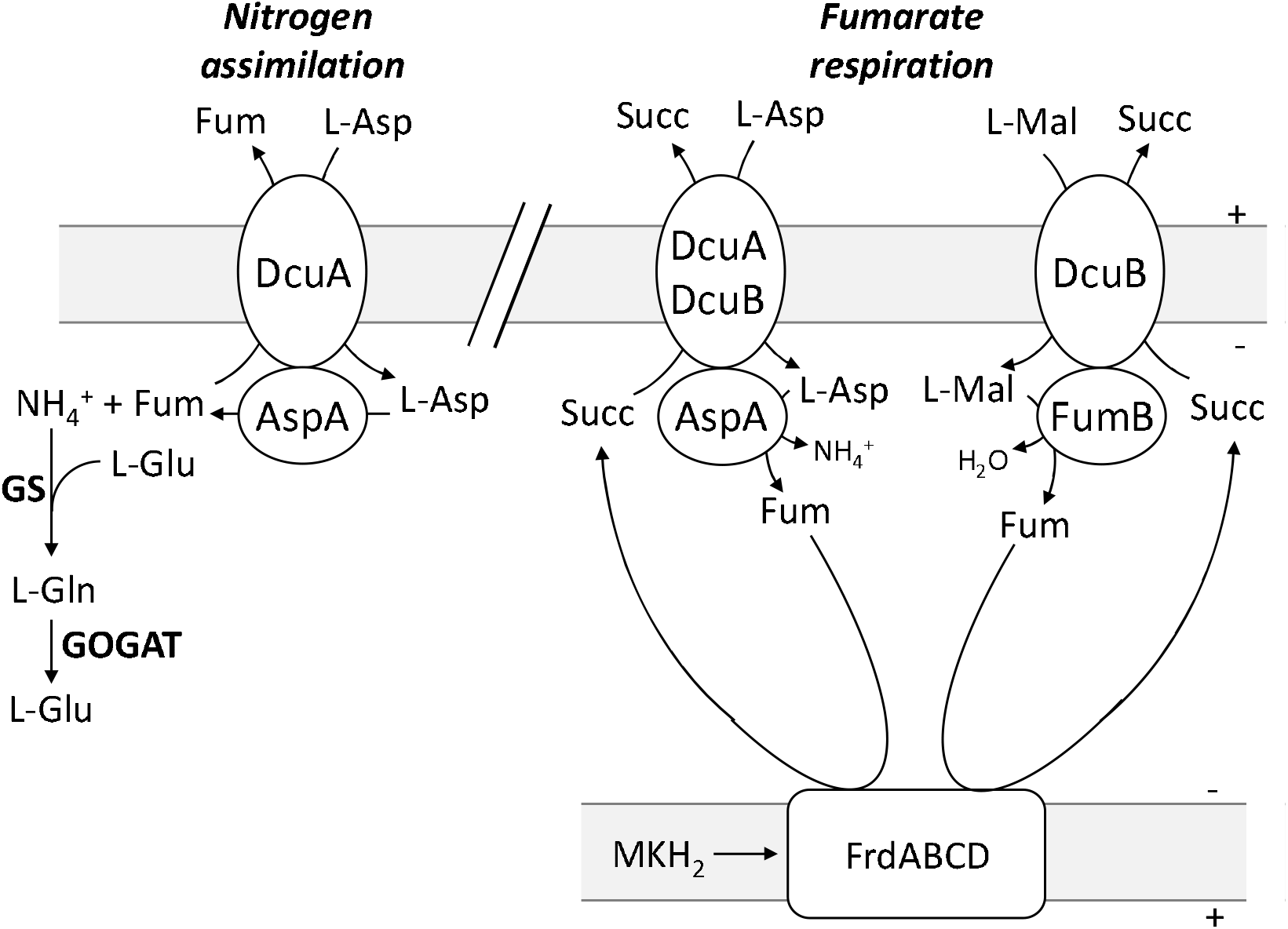
Tentative scheme for the DcuA/AspA, DcuB/AspA and DcuB/FumB complexes and the coordinated reactions in fumarate respiration and nitrogen assimilation from L-Asp. Abbreviations as in Fig. 1.

When L-Asp functions as the nitrogen source for bacterial growth, it can be used as the sole source of nitrogen (Strecker *et al.* 2018). The ammonia released by AspA feeds the route for nitrogen assimilation via glutamine synthetase GS and glutamine 2-oxoglutarate amino transferase GOGAT (Strecker *et al.* 2018). The deaminating activity of AspA is further stimulated by the regulator GlnB (or PII), which responds to the nitrogen status of the bacteria (Schubert *et al.* 2020). In consequence, from the L-Asp only the ammonia used under conditions of nitrogen assimilation, and fumarate with some L-malate is released by DcuA in near stoichiometric amounts to L-aspartate uptake (Strecker *et al.* 2018). Thus, coordination of the pathway with DcuA and AspA in the DcuA/AspA metabolon (Fig. 4B) is also highly indicated by genetic colocalization of the genes, protein interaction and the coordination of the L-Asp/Fum antiport to the ammonia supply.

## Materials and Methods

### Bacterial strains and growth conditions

The *E. coli* K12 strains and the plasmids are listed in Table 1. All molecular methods, including cloning, DNA isolation, and manipulations were performed according to standard procedures (Jones and Gunsalus 1987; Sambrook *et al.* 2001; Miller 1992; Zientz *et al.* 1998; Kleefeld *et al.* 2009a). The oligonucleotide primers are shown in Table 2. Bacteria were grown aerobically and anaerobically at 37°C in lysogeny broth (LB).

**Table 1.** Strains of *E. coli* and plasmids used.

| Strain | Genotype | Reference |
| --- | --- | --- |
| C43(DE3) | Mutant of BL21(DE3), for expression of membrane proteins | Miroux and Walker 1996 |
| BTH101 | F- *cya-99, araD139, galE15, galK16, rpsL1 (Strr), hsdR2, mcrA1, mcrB1* | Karimova *et al.* 1998 |
| Plasmid | Genotype | Reference |
| pKT25 | C-Terminal T25 protein fusion plasmid, pSU40 derivative, KanR | Karimova *et al.* 2001 |
| pKNT25 | N-Terminal T25 protein fusion plasmid, pSU40 derivative, KanR | Karimova *et al.* 2001 |
| pUT18 | N-Terminal T18 protein fusion plasmid, pUC19 derivative, AmpR | Karimova *et al.* 2001 |
| pUT18C | C-Terminal T18 protein fusion plasmid, pUC19 derivative, AmpR | Karimova *et al.* 2001 |
| pKT25-Zip | T<sub>25</sub>Zip expression plasmid, pKT25 derivative, KanR | Karimova *et al.* 1998 |
| pUT18C-Zip | T<sub>18</sub>Zip expression plasmid, pUT18C derivative, AmpR | Karimova *et al.* 1998 |
| pMW2367 | AspA<sub>T18</sub> expression plasmid, pUT18 derivative, AmpR | This study |
| pMW2368 | T<sub>18</sub>AspA expression plasmid, pUT18C derivative, AmpR | This study |
| pMW3086 | FumB<sub>T18</sub> expression plasmid, pUT18 derivative, AmpR | This study |
| pMW2376 | Expression of DcuC<sub>1-229</sub>-T25-DcuC<sub>230-461</sub>, ('DcuC<sub>SWT25</sub>') pKNT25 derivative, KanR | Strecker 2018 |
| pMW1028 | Expression of DcuB1-211-T25-213-446 (,DcuB<sub>SWT25</sub>'), pKNT25 derivative, KanR | Wörner *et al.* 2016 |
| pASK-IBA3plus | Expression plasmid with tet promoter and C-terminal Strep-tag, AmpR | IBA, Göttingen |
| pMW3049 | pASK-IBA3plus with *aspA*-Strep, AmpR | Schubert *et al.* 2020 |
| pBAD18 | Expression vector; pBR322 *ori*, arabinose-inducible pBAD promoter, KanR | Guzman *et al.* 1995 |
| pMW577 | pBAD18 with *dcuA-phoA*, KanR | Bauer *et al.* 2011 |

**Table 2:**
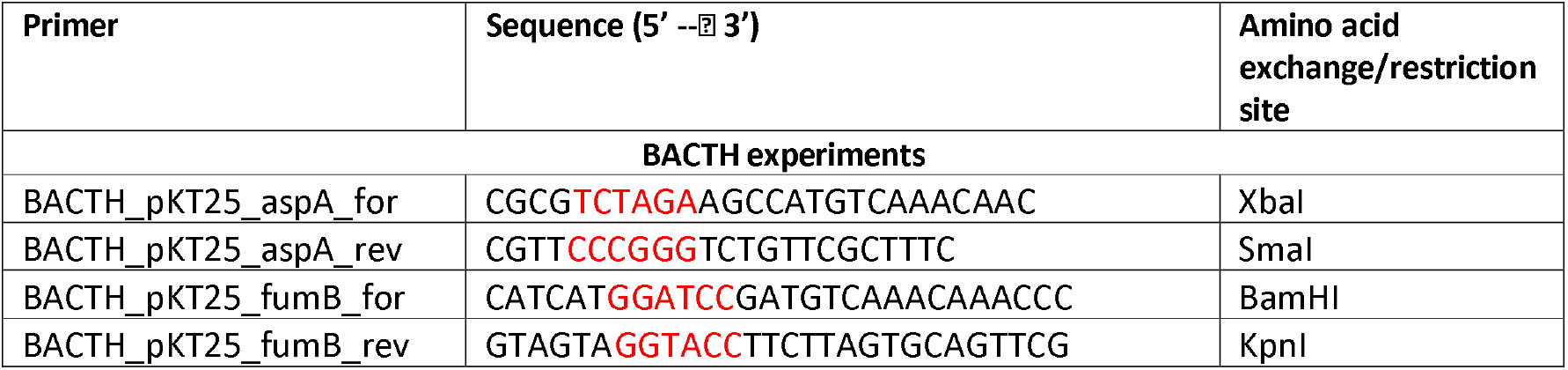
Oligonucleotide primers. Primer for PCR amplification for AspA, FumB and EIIA. Restriction sites and amino acid exchange positions are printed in red.

### β-Galactosidase assay

Cells for the BACTH measurements were grown anaerobically in LB medium. Interactions in the BACTH system were measured in terms of the β-galactosidase activity (Miller 1992). BACTH experiments were conducted as described previously (Monzel *et al.* 2013) with slight modifications (Wörner *et al.* 2016). Activities are the mean of at least two independent experiments and four replicates each.

### Membrane protein interaction (mSPINE)

The mSPINE assay for protein interactions was performed as described previously (Graf *et al.* 2014; Wörner *et al.* 2016). *E. coli* strain C43 was co-transformed with plasmids, pMW3049 and pMW577, which encoded the fusion proteins AspA-Strep and DcuA-PhoA, respectively. The bacteria were grown aerobically in 400 ml LB medium to an OD_578_ of 0.5. Protein expression was induced for 3 h with 100 μM L-arabinose and 100 μM anhydrotetracycline (AHT). To crosslink the proteins *in vivo,* the bacteria were incubated with formaldehyde (0.6% w/v) for 15 min at 37°C with shaking. Next, the bacteria were harvested with centrifugation (11300 × *g* for 10 min), and the sediment was washed in Tris-HCl buffer (pH 8, 50 mM). For Strep-Tactin purification, cells were disrupted with a French Press (1260 psi) in a buffer that contained lauryldimethylamine oxide (LDAO,0.05% (v/v)). The cell homogenates were clarified by centrifugation (39100 × *g* for 30 min at 4°C) and subjected to Strep-Tactin chromatography, 10% SDS-PAGE, and Western blotting. Western blots were probed with anti-Strep and anti-PhoA antisera (IBA Lifesciences, Sigma-Aldrich) (Scheu *et al.* 2010; Graf *et al.* 2014; Wörner *et al.* 2016).

### SDS-PAGE, Western Blot, and Immunostaining

SDS-PAGE and Western blots were performed according to published procedures (Graf *et al.* 2014; Wörner *et al.* 2016). Immunostaining was performed with horse-radish peroxidase (HRP)-coupled anti-His-HRP, anti-Strep-HRP, and anti-IgG-mouse-HRP polyclonal antiserum (Sigma-Aldrich) or with anti-PhoA antibodies (produced in mouse, Sigma-Aldrich). For visualization chemiluminescent substrate HRP (Merck Millipore) was used and the blots exposed on X-ray film (Advansta).

## Acknowledgments

We are grateful to Dr. A. Strecker (Mainz) for preparing the plasmids with the DcuA_SWT25_ and DcuC_SWT25_ constructs. Financial support from Deutsche Forschungsgemeinschaft (grant UN 49/19-1) is gratefully acknowledged.

